# TSCDA: A novel greedy approach for community discovery in networks

**DOI:** 10.1101/2021.10.08.463718

**Authors:** Arman Ferdowsi, Alireza Khanteymoori, Maryam Dehghan Chenary

## Abstract

In this paper, we introduce a new approach for detecting community structures in networks. The approach is subject to modifying one of the connectivity-based community quality functions based on considering the impact that each community’s most influential node has on the other vertices. Utilizing the proposed quality measure, we devise an algorithm that aims to detect high-quality communities of a given network based on two stages: finding a promising initial solution using greedy methods and then refining the solutions in a local search manner.

The performance of our algorithm has been evaluated on some standard real-world networks as well as on some artificial networks. The experimental results of the algorithm are reported and compared with several state-of-the-art algorithms. The experiments show that our approach is competitive with the other well-known techniques in the literature and even outperforms them. This approach can be used as a new community detection method in network analysis.

## 1 Introduction

One of the fundamental topics in the study of networks that has attracted much attention recently is partitioning nodes into so-called communities consisting of highly interactive vertices. In this sense, a network can be modeled as a graph *G* = (*V, E*) with the set of vertices *V* and edges *E*. Despite there is no universally agreed-upon definition of a community in the literature, this work uses one of the most common ones: a community is a subset of vertices *C* ⊆ *V* with a high density of edges between nodes inside the subset and a low density of edges connecting this subset to the others. Accordingly, the community detection problem is the problem of finding a partitioning of the nodes into communities.

Such a partitioning indded enables us to survey many underlying properties of networks and has numerous applications in a wide range of areas including WEB [4], social media/network analysis [33], [59], [2], biological networks [25], image segmentation [38], pattern recognition [23], data clustering [40], recommender systems [3], and etc.

There are several variants of the problem depending on whether: the network is weighted or unweighted, the network is directed or indirected, the desired number of communities is given or not, and communities have overlap or they are completely separated; e.g., see [27], [13], [9], [53], [17], [42]. On the other hand, various approaches for discovering communities have been proposed, and among them, there is a wide range of non-optimization-based algorithms such as the Label Propagation algorithm [48]. However, the majority of community detection approaches follow a simple optimization strategy: they first define or pick a specific function for determining the quality of communities which is called quality function, and then they come up with an optimization algorithm that can be used to find communities with respect to optimizing the value of the quality function. In the following, we first give an overview of some of the most well-known quality functions and some algorithmic approaches for discovering communities.

In the literature, there are several quality measures to qualify the goodness and accuracy of partitioning. At a glance, according to [11], community quality functions can be classified into four broad categories: internal-connectivity-based metrics (IC), external-connectivity-based metrics (EC), metrics based on both internal and external connectivity (I/EC), and metrics based on networks’ topology.Table 1 lists some of the well-known connectivity-based quality functions. On the other hand, among the metrics based on the topology of the network, *Modularity*, introduced by Newman [43], is one of the most used and best-known quality functions. For a community *C*, Modularity is defined as the number of edges within *C* minus the expected number of such edges. In addition to all this, there are plenty of other community score functions such as Flake-ODF, Cohesiveness, Clustering coefficient, Spart, Permanence, etc. We refer the reader to [11], [16] for a comprehensive survey about quality functions for community analysis.

**Table 1:**
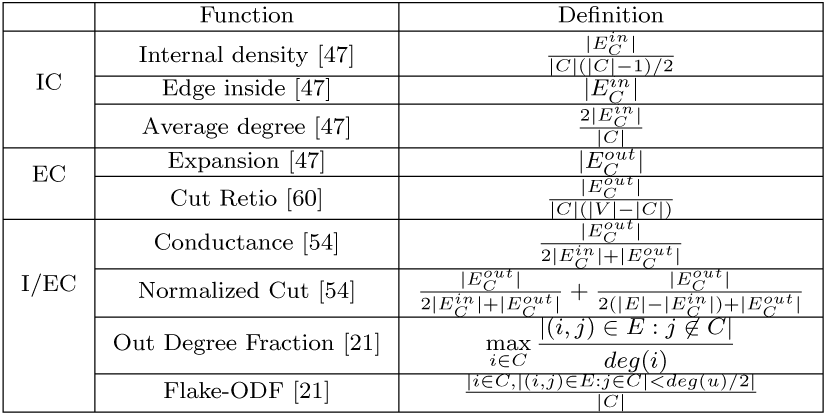
Several connectivity-based quality functions for measuring the quality of a community *C*. 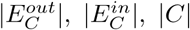 and *deg*(*i*) respectively represent the number of external connections of *C*, number of internal connections of *C*, the number of vertices belonging to *C*, and the degree of node *i*.

Naturally, devising and considering a quality function is expected to be followed by proposing an algorithmic technique for discovering communities with respect to optimizing the quality function. There have been relatively few exact and explicit approaches, such as using linear or integer programming techniques (see [1]). Even though these methods can obtain very accurate results, they usually fail to encounter extensive networks since optimizing the quality functions often falls into the category of difficult computational problems [50]. Therefore, in most cases, a heuristic method need to be inevitably employed to increase the (time) efficiency [20]. Accordingly, while dealing with massive networks, which we usually face in the real world, one acceptable approach would be to rely on heuristic techniques from the very beginning [12]. Indeed, many heuristic (optimization-based) algorithms tackle the community detection problem through one or a combination of the following approaches:

– Bottom-up: starting with singleton communities and merging communities.
– Top-down: starting with considering one big community and splitting clusters.
– Local search: starting with an initial partition and migrating nodes.

For instance, Blondel et al. [8] proposed a bottom-up heuristic greedy Modularity maximization algorithm named *Louvain. Infomap* algorithm [49] is another bottom-up algorithm and uses the same optimization procedure as in the Louvain algorithm but tries to optimize a different quality function, an entropy-based function called *Map Equation. Fast Greedy* algorithm, which was developed by Newman [42], uses a greedy hierarchical bottom-up approach. *Girvan-Newman (GN)* algorithm is, on the other hand, a top-down hierarchical method based on betweenness metrics [27]. This is while, the spectral graph partitioning approach discovers communities of a network with respect to solving relaxed versions of the *Normalized cut* minimization problem. This approach uses information from the eigenvalues (spectrum) of special matrices associated with a graph and uses the corresponding eigenvectors to detect communities; see [58], [24]. To summarize, for giving a better vision to some recent progress in community detection algorithms, we refer the reader to [12], [35], [31], [57].

The growing trend in defining and developing different quality functions can rightly indicate that all of these function have their own weakness in applications. For example, Modularity, as one of the mostly-used measures in this area, suffers from several known drawbacks such as the resolution limit (i.e., failure to detect communities smaller than a specific scale) [22], and the high degeneracy (i.e., leading to various partitions with equally high Modularity score) [28]. On the other hand, one downside of many connectivity-based quality functions is that they usually only consider the number of connections between nodes and do not simultaneously consider other topological features of the network, such as distance between vertices, that may lead to obtaining almost poor results. Furthermore, another argument is that regardless of which specific measure needs to be optimized, many optimization algorithms lack in their nature. For instance, many approaches start with relatively inaccurate initial communities that makes it difficult, impossible, or even very time-consuming to reach the final high-quality communities [61], [51].

### Main contribution

This paper aims to discover a set of non-overlapping communities ℂ = {*C*_1_, …, *C*_*k*_} for a given unweighted undirected network, where the number of communities, *k*, will be automatically estimated by the method. To do so, we first modify one of the fundamental connectivity-based quality functions. Not only do we take the network connectivity into consideration, but we also investigate the role of the network’s influential nodes in establishing communities. Further, despite the fact that this minor modification can alone cause to obtaining better communities, we also propose an efficient optimization-based algorithm to detect high-quality communities with respect to optimizing the modified quality measure. The algorithm greedily establishes high-quality initial communities based on recognizing the influential vertices, which are at an approximate proper distance from each other. It then iteratively updates the obtained communities to reach final communities with the best possible quality value in a local search manner.

For the evaluation part, we apply our proposed algorithm and also five state-of-the-art heuristic community detection algorithms on several well-known real-world networks as well a famous category of randomly generated graphs whose optimal community structures are available. We then compare the obtained communities with each other. The results demonstrate the superiority of our proposed algorithm over the other under-study algorithms.

The rest of the paper is organized as follows. First, we introduce the community quality function and the community mining algorithm in Section 2, and in Section 3, we report the experimental results.

## 2 TSCDA: Two-Stage Community Detection Algorithm

In what follows, we first set up some notations and preliminaries, which we use throughout the paper. After that, we recommend our quality function and then, we propose our community detection algorithm.

### 2.1 Background and notations

Let *G* = (*V, E*) be a network with *n* = |*V* | nodes, *m* = |*E*| edges, and the adjacency matrix *A*. Let ℂ = {*C*_1_, …, *C*_*k*_} denotes a set of *k* disjoint subsets of *V*. We refer to each *C*_*l*_ ∈ ℂ as a community of *G*. The followings are more notations:

– For each *i* ∈ *V*, let *deg*(*i*) shows the degree of *i*.
– Let *N*_*G*_(*i*) = {*j* ∈ *V* | (*i, j*) ∈ *E*} denotes the set of neighbors of *i*.
– For each *C* ∈ ℂ, we define: 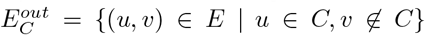 and 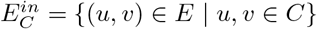 to respectively be the set of *outer* and *inner edges* of *C*.
– A vertex *i* ∈ *V* is said to be ℂ−*homeless* if it does not belong to any *ℂ* ∈ ℂ. The set of all ℂ−*homeless* vertices is denoted by *H*_ℂ_. Note that *H*_ℂ_ = *V* when ℂ = ∅.
– For every vertex *i* ∈ *V*, let *C*(*i*) ∈ ℂ be the community that contains *i*.
– For every *i* ∈ *V*, we let 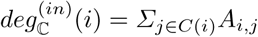 and 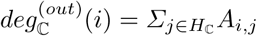 to be the ℂ−*inner degree* and ℂ−*outer degree* of *i*, respectively. Note that 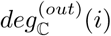 is the number of edges connecting node *i* to the homeless vertices.
– Distance between any two vertices *i, j* ∈*V* is defined by the length of the shortest path connecting these two nodes in *G*, and is denoted by *dist*(*i, j*). The diameter of *G* is the longest shortest path length in the graph and denoted by *diam*(*G*).
– Let *S* ⊂ *V*. For any *α* ∈ {2, 3, …, *diam*(*G*)}, vertex *i* ∈ *V* − *S* is said to be *α-far* from *S* if 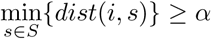. Note that every vertex *i* ∈ *V* is *α-far from S* when *S* = ∅.
– Two communities *C* and *C*^′^ are said to be *neighbors* if there exists an edge (*i, j*) ∈*E* connecting *i* ∈*C* to *j* ∈*C* ′ In this case, we refer to both *i* and *j* as *boundary vertices* of *C* resp. *C*^′^. Let *BV* (*C*) denotes the set of all boundary vertices of *C*.

### 2.2 The quality function

Without any doubt, the degree of a vertex can usually be thought of as one of the authority factors of the vertex for controlling/affecting/biasing its surrounding nodes. In fact, high degree vertices can impose many features on and pass various messages to their nearby vertices. They can even influence and bias their surrounding nodes’ attributes and behaviors [30], [62], [34], [46], [10]. To be more specific, a large number of (complex and social) networks usually contain a proportionally small number of influential high-degree vertices e.g., celebrities in a social media or essential and top-selling commodities in a copurchasing network, which are surrounded and followed by a vast number of lower degree vertices; see [45], [18], [7], [14]. In such networks, each vertex is either connected directly to or is usually only a few links away from a high-degree node. Consequently, high-quality communities are likely expected to be formed around high-degree vertices. Indeed, we have observed that high-performance communities typically contain one or a few numbers of high-degree vertices, along with a large number of their nearby surrounded lower-degree vertices.

In this vein, we refer to a vertex *i* ∈*C* ∈ ℂ with the maximum ℂ −inner degree as an *influential node* of the community *C* and denote it by *m*(*C*). Note that, in the present paper, every community *C* ∈ ℂ has only one influential node; if there are more than one vertex in *C* with the same maximum inner degree, one of them will be selected arbitrarily. Based on the above consideration, we define the distance between a vertex *i* and a community *C* to be *dist*(*i, m*(*C*)). In addition, *dist*(*m*(*C*), *m*(*C*^′^)) represents the distance between two communities *C* and *C*^′^. We also define the *radius* of a community *C* ∈ ℂ to be 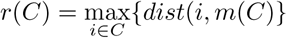.

Furthermore, we have also noticed that an influential node’s effect on other vertices is somehow directly related to the distance between them. It means that, for example, a very famous individual in a social network can have a more significant impact on people who follow him/her directly (distance= 1) rather than those who know him/her through other people (distance *>* 1) (see [68] for example). Therefore, we conjectured that a community in which the overall distance between the influential node and the other vertices is small (i.e., its radius *r* being small) is likely to be a more interactive and homogeneous community.

Accordingly, we obtain the quality of a community *C* ∈ ℂ as follows:

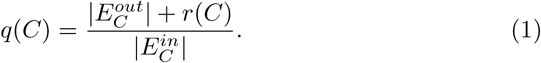

We call *q*(*C*) the *quality value* of the community *C*. Besides, let 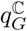 indicates the quality of the partition ℂ of *G* and be defined as follows:

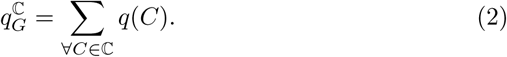

Note that the lower the value of *q*(*C*), the higher the quality of the community *C* is. Basically, the proposed community quality measure determines the quality of a community *C ∈ℂ*by adding up the following two terms: 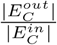 and 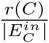. Recall the definition we presented in Section 1 for a community. Based on that, it is natural to think that a high-quality community has a large number of outer edges and a small number of inner edges at the same time. In other words, 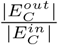 measures the independence of the community *C* from the rest network, and it becomes small (close to zero) for a community with a high density of edges between its vertices and a low density of edges connecting its vertices to the other communities.

For a better understanding, let us consider the following case. Assume that *C*_1_ and *C*_2_, shown in Figure 1, are two communities of a network with influential vertices *a* and *b*, respectively. Consider, for example, the three quality functions 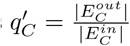, conductance 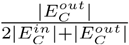, and modularity. The score of each of these quality functions for *C*_1_ and *C*_2_ are equal, and therefore based on their viewpoint, *C*_1_ and *C*_2_ have the same quality. This is while it is apparent that *C*_2_ is a much worse community due to its more absence links than *C*_1_. Nevertheless, if we consider our proposed quality function, we have 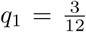 and 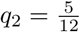, which truly shows that *C*_1_ is a better community.

**Fig. 1:**
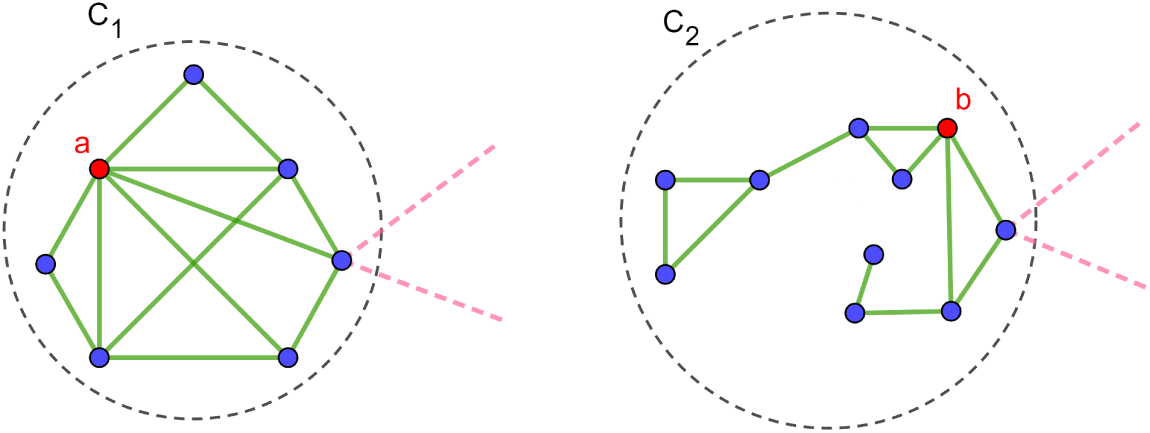
Two communities *C*_1_ and *C*_2_ with influential vertices *a* and *b*, respectively. *C*_2_ is a worse community since it has more disconnected node pairs than *C*_1_. 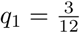 and 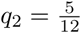 demonstrate the superiority of *C*_1_ over *C*_2_.

### 2.3 The algorithm

In this section, we are going to depict our Two-Stage Community Detection Algorithm (TSCDA). For a given simple network *G* = (*V, E*), the algorithm tries to find a set of communities ℂ = {*C*_1_, *…, C*_*k*_} for which 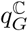 is (approximately) minimized. TSCDA consists of two phases: *Creating Initial Communities* and *Updating Communities*, which we describe as follows:

#### Stage 1: Creating Initial Communities (CIC)

Algorithm 1 elaborates on the PseudoCode of this stage. This phase aims at establishing a set of initial communities by guessing promising influential vertices. Recall the definition of the influential nodes provided in Section 2.2. Actually, it can be easily turned out that as important as the proximity of each community’s vertices to the corresponding influential node is as crucial as separating communities. Indeed, considering an adequate approximate distance between initial communities can lead to a significant breakthrough for finding final accurate communities. This is in accordance with the fact that discovering high-quality communities relies considerably on recognizing initial influential nodes, which are at a proper distance from each other. As a result, here for establishing initial communities, we put an extra condition on defining the influential nodes: we introduce a parameter *α* to supervise the distance between influential vertices.

Basically, for a given parameter *α* ∈{2, 3, …, *diam*(*G*)}, this step guesses a set of initial influential nodes *M* (*α*). Then, it determines a set ℂ(*α*) consisting of communities, each of which is associated with an influential vertex. The following steps present the procedure.

– Initialize *M* (*α*) = ℂ(*α*) = ∅.
– While there is an *α*−far vertex from *M* (*α*), repeat the following operation:
  – Among all *α*−far vertices from *M* (*α*), select a ℂ(*α*)−homeless node *i* with a maximum ℂ(*α*)−outer degree ^1^ as a new influential vertex and assign it into both of the sets *M* (*α*) and C(*α*).
– Assign every ℂ(*α*) homeless node to its closest community.
– Update the communities’ influential nodes (resp. *M* (*α*)) based on the definition provided in Section 2.2, if necessary.

The above procedure will be repeated for every *α* ∈{2, 3, …, *diam*(*G*) ^2^}, and among all the obtained sets of communities C(*α*), one with the least 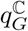 will be selected. By doing this, we will manage to find an appropriate between-communities distance with a good approximation. We refer to the best obtained C(*α*) as the set of initial communities. Besides, it should be noted that obtaining C(*α*) for different values of *α* does not unacceptably increase the execution time since almost many (social and complex) networks are sparse and have a relatively small diameter compared to their number of vertices; See [52], [4], [14], [7], [44].

##### Algorithm 1

Creating Initial Communities (CIC)

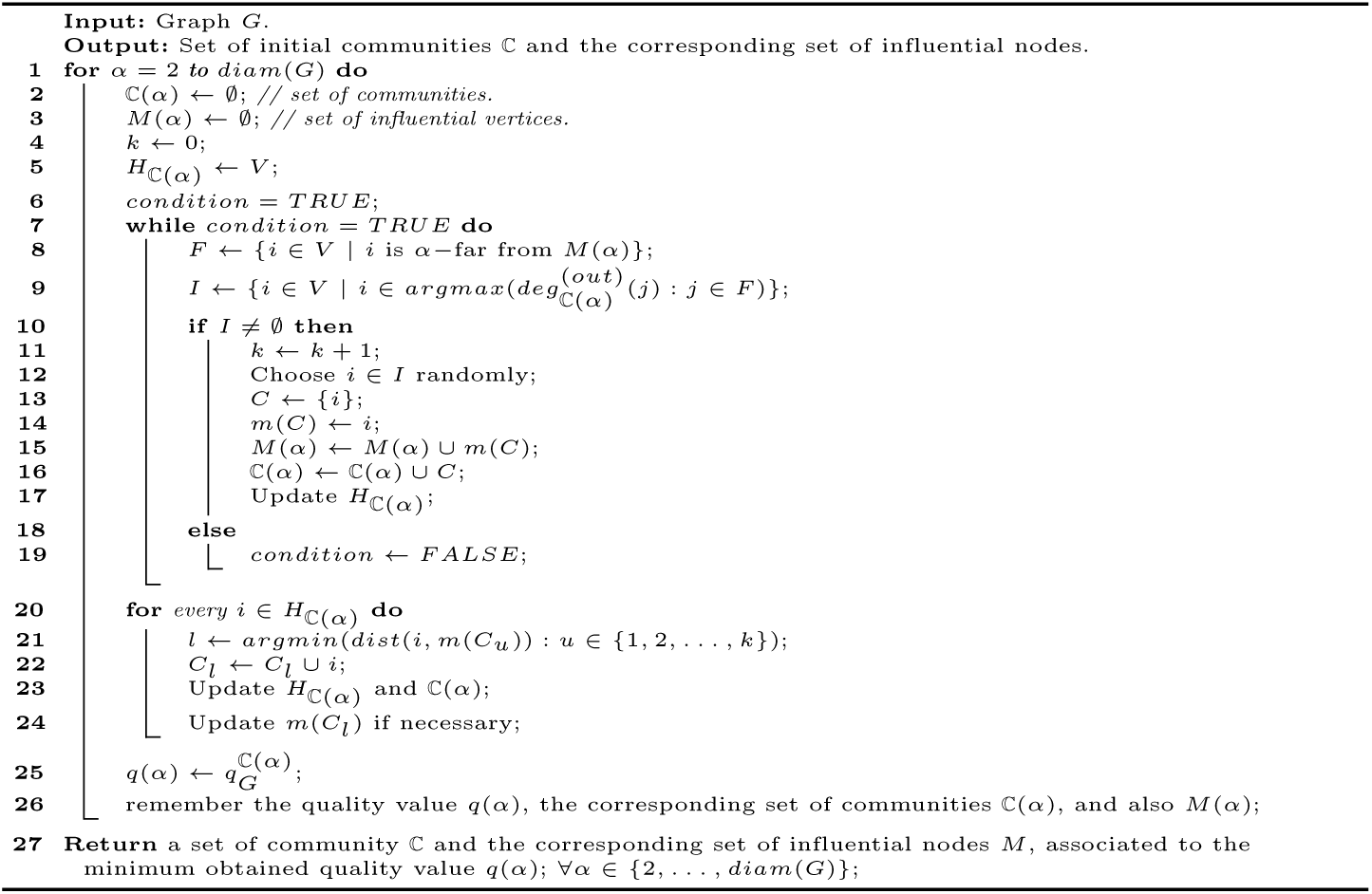

### Stage 2: Updating Communities (UP)

Algorithm 2 provides the PseudoCode of this stage. This phase basically updates the initial communities by using a local search technique bases on iteratively moving to neighbor solutions. In a more precise explanation, it iteratively selects a community in such a way that the probability of choosing a community *C* with a worse quality value is more than a community with a better quality value. After that, it greedily improves the communities based on one of the three functions *Merge, Split*, and *Swap* at a time. These functions operate as follows:

– *Merge* function checks if merging *C* into any of its neighbor communities can improve the quality value.
– *Split* function checks if *C* can be divided into more than one community, given that the quality value of the obtained division is better than *q*(*C*), the quality value of *C*. To do so, the CIC stage (Algorithm 1) will be applied to the induced sub-graph *G*^′^ of *G* formed from the subset *C* of *V*.
– *Swap* function applies the best possible swaps between nodes in *C*, and nodes belong to *C*’s neighbor communities.

After each iteration, the best function that leads to obtaining the most significant improvement in the quality value will be selected. The communities and the influential vertices will be updated afterward. The procedure terminates when no improving local search move exists.

It is apparent that the better the obtained initial communities are, the less number of reforms is needed to be done by the updating community (UC) stage. In the next section, we will discuss this argument in detail. More specifically, we will show how these two stages, individually resp. together, lead to obtaining high-quality communities.

## 3 Experiments

Regardless of optimizing the quality function considered, the ultimate target of a community detection algorithm is to detect high-quality communities, which means finding communities that are as similar as possible to the optimal community structures, of course, if such structures are available. To be more precise, according to [11], [56], obtaining communities that leading to a near-optimal value of quality functions might bear no similarity to the optimal community structures. Thus even by discovering communities with optimal quality function, other verification methods are needed to confirm the efficiency of the communities. In this part, we aim to evaluate our algorithm’s performance against several most successful community detection algorithms. To make a consistent condition for the comparisons, we try to estimate the similarity between each algorithm’s discovered communities and the optimal community structures. Therefore, in this work, we take networks into account whose optimal community structures (i.e., *ground truth*) are available in the literature. We consider six well-known real-world datasets as well as a commonly-used class of synthetic graphs, and to determine the similarity between each algorithm’s obtained communities and the ground truth, we use two different evaluation metrics (explained in Section 3.2).

### Algorithm 2

Updating Communities (UC)

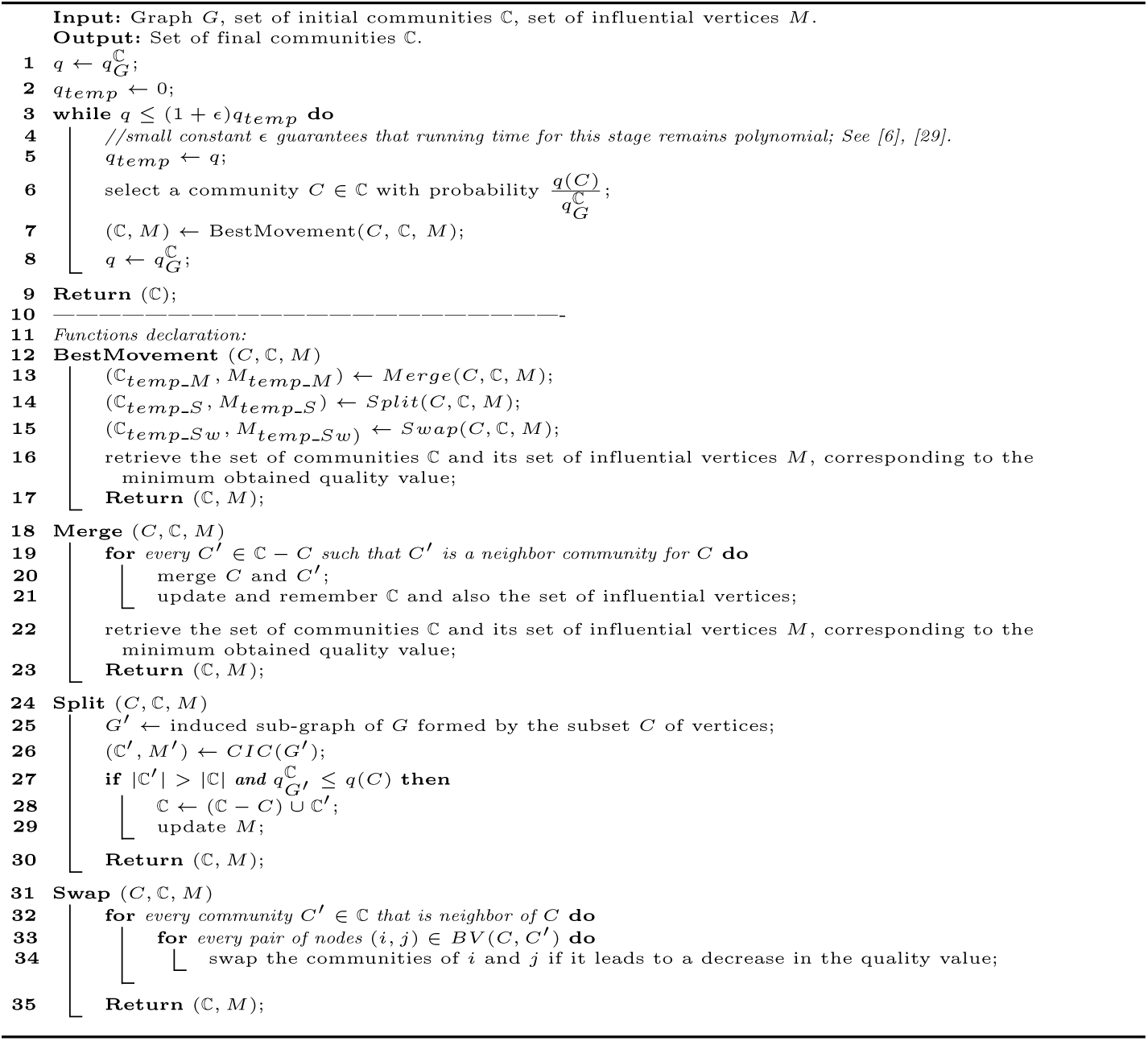

All tests are conducted on a computer system with a processor Intel(R) Core(TM) i5-7300*U* CPU @ 2.60GHz, 2712 Mhz, 2 Core(s), 4 Logical Processor(s), 8 GB of Rams, and Win10 OS. Moreover, algorithms are implemented by the C++ programming language.

### 3.1 Datasets

The examined real-world networks in this paper have various properties and characteristics and are listed in Table 2. In addition, as synthetic benchmarks, we use the LFR artificial networks [36] that highly resemble real-world social networks and have a prior known and built-in community structure as well. The generating process of such artificial graphs is based on several user-defined parameters and is fully explained in [36]. These networks consider power-law distributions for both the degree and the community size with exponents *γ* and *β*, respectively. In addition, The mixing parameter *μ* specifies that every vertex of the graph should share a fraction of 1 − *μ* of its links with the other nodes of its community and a fraction of *μ* with the other nodes of the network. *μ* can control the noise, and the more the value of *μ*, the worse the community structure emerges (i.e., communities get less distinct). Therefore, for a considerable weight of *μ*, community detection algorithms face more trouble finding the optimal communities [19]. In this work, we use four different LFR graphs with four different mixing parameters *μ* (*μ* = 0.1, 0.3, 0.6, and 0.9), each of which has 1000 nodes and 5 communities. All the other parameters are set to the default values explained in [36].

**Table 2:** Real-world networks

| Network | $n$ | $m$ | number of communities | diam |
| --- | --- | --- | --- | --- |
| Karate club [67] | 34 | 78 | 2 | 5 |
| Dolphin network [39] | 62 | 159 | 2 | 8 |
| American college football [27] | 105 | 613 | 12 | 4 |
| Email-Eu-core [37] | 986 | 16064 | 42 | 7 |
| Amazon network [41], [65] | 334863 | 925872 | 49732 | 44 |
| LiveJournal social network [55], [65] | 3997962 | 34681189 | 287512 | 17 |

### 3.2 Evaluation metrics

As mentioned earlier, once the communities are discovered, the next job might be to estimate the detected structures. The evaluation becomes more reliable if the ground truth is available. It is natural to conjecture that if the algorithm can detect a community structure with a high resemblance with the ground truth, it is trustworthy enough to be used further for other networks for which the underlying optimal community structures might not be available. Consequently, various metrics, such as F-measure [5] and Rand index [32], have been developed to correspond between the detected communities and the ground truth. Nevertheless, each of them has some shortcomings. For instance, it turned out that F-measure does not reveal the mutual information among communities, while the Rand index suffers from having biases that might lead to low-quality results. As a result, most of the research works considerably rely on two robust performance metrics named *Adjusted Rand Index (ARI)* and the *Normalized Mutual Information (NMI)*. In the following, we introduce these two quantitative metrics for validating the output of a community detection algorithm. ARl can be determined easily, and it measures the extent to which the two partitions agree with one another by evaluating the relationship between pairs of nodes. However, NMI is more complicated to compute but can quantify the performance of different community structures with high computational complexity.

Suppose that for a given network *G*, ℂ = {*C*_1_, …, *C*_*k*_} and 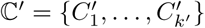 be respectively a set of communities obtained by an algorithm *A* and the ground truth.

Normalized Mutual Information (NMI) [15] is indeed a well-known clustering comparison metric and can be used to evaluate the similarity between the optimal communities and the discovered ones. The NMI value corresponding to the algorithm 𝒜 can be written as

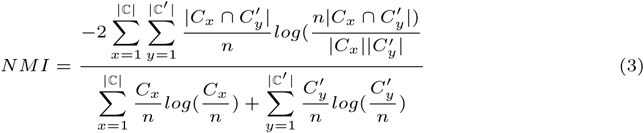

If the detected communities are identical to the ground truth, the NMI takes its maximum value one, while in the case where the two sets totally disagree, the NMI score is zero. Generally, the more the NMI, the better community structures have been found.

Adjusted rand index (ARI) [66], corresponding to the algorithm *A*, measure the similarity between ℂ and ℂ′as follows

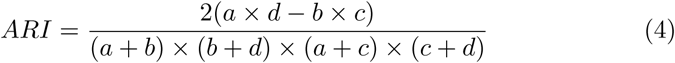

where *a, b, c* and *d* are respectively the number of vertex pairs that are in the same community in both ℂ′ and ℂ, in the same community in ℂ′but not in ℂ, in the same community in ℂ but not in ℂ′, and in different communities in both ℂ′ and ℂ. Like the NMI measure, the value of *ARI* varies between 0 and 1, and the higher it is, the more similar the communities obtained by the algorithm 𝒜 are with the grand truth of *G*.

### 3.3 Comparison algorithms

Recall that one of the most widely used quality functions for community detection is Modularity. The reliance of many of the community discovery methods on this quality function and their success in many social and biological networks led us to measure our algorithm’s performance against three state-of-the-art Modularity maximization algorithms, the Distributted Louvain (*DLouvain*) [26], Simulated Annealing [63], and FastGreedy [42] algorithms. Besides, we also consider two other high-efficient algorithms: Infomap [49], and the Improved Label Propagation [64]. These two algorithms are respectively based on (i) optimizing a well-known information-theoretical measure (description code length quality function), which returns high-quality communities for big networks with a relatively low level of noise, and (ii) introducing a new similarity measure and using the minimum distance and local centrality to establish initial communities. The algorithm then employs the label influence to update the community structures based on the new similarity measure.

It’s worth mentioning that all these algorithms are among the best high-efficient and powerful community detection algorithms for social and complex networks [56].

### 3.4 Experiment setup and results

For arranging the comparisons over real-world networks, we compare the performance of all introduced algorithms by examining the average ARI and NMI ranks over all datasets. Figure 2 presents the results. One can conclude that our method significantly outperforms other algorithms, i.e., the communities detected by the TSCDA are much more like the ground truths than those of the other methods. In addition, Figure 3 illustrates the ARI and NMI values obtained by applying TSCDA and other comparable algorithms to the LFR benchmarks explained in Section 3.1. As we can see, although the performance of all the algorithms decreases with increasing *μ*, TSCDA surpasses other techniques in all cases.

**Fig. 2:**
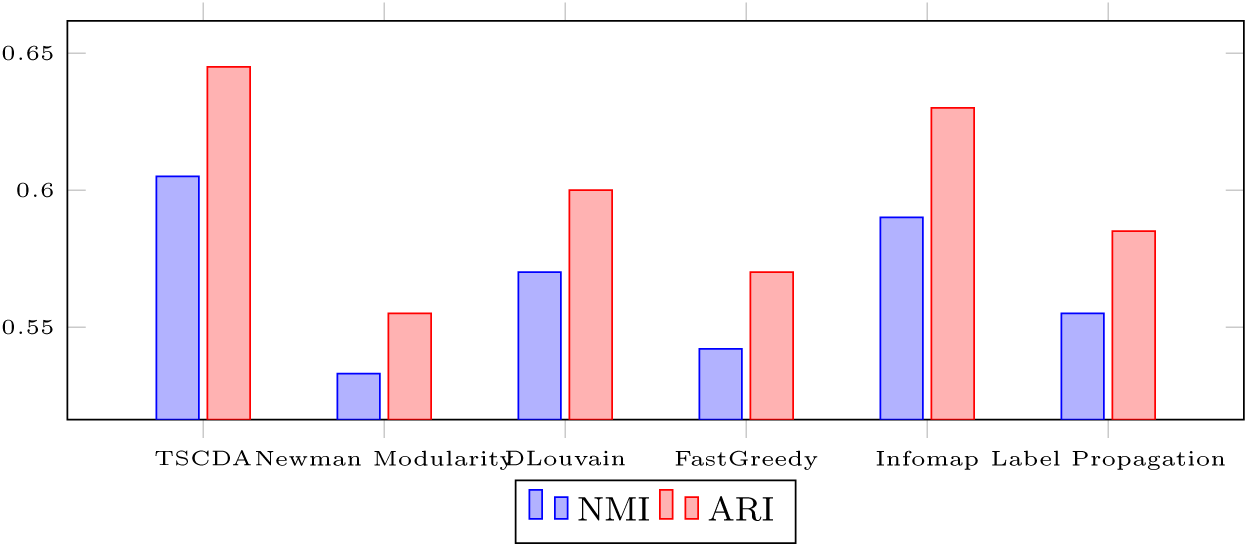
Average normalized ARI and NMI performance rank of different algorithms applying to the real-world networks, presented in Table 2. The average NMI and ARI values obtained by TSCDA are respectively equal to 0.605 and 0.645.

**Fig. 3:**
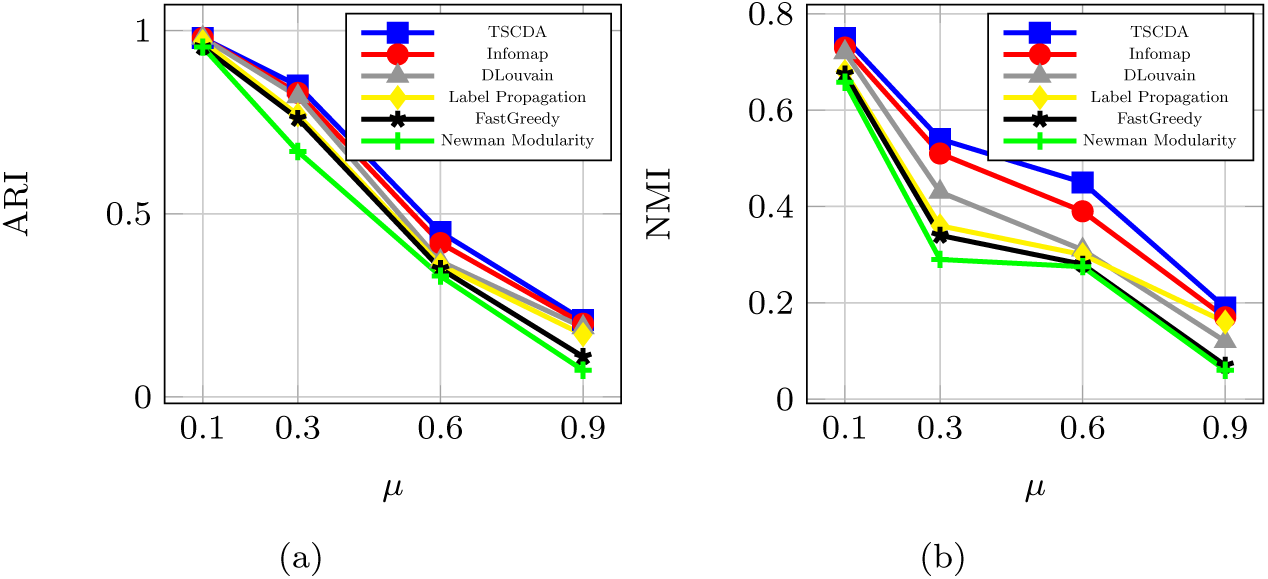
ARI (a) and NMI (b) performance rank of each algorithm applying to the LFR synthetic graphs, generated by considering four different mixing parameters *μ*.

Another important argument is that it would be substantial to see how accurate the first stage of the TSCDA works. In other words, we have to check how authoritative the obtained initial communities are and how many reforms must be made by the second stage. Table 3 provides a comprehensive comparison of these two steps based on all the networks studied. In addition to the ARI and NMI values, obtained by each stage, the following information is also depicted in the table: the best values for *α* founded by the CIP stage, the number of iterations done by the UC stage, the number of communities estimated by each of the stages, and finally, the amount of execution time elapsed by each stage. By noticing the ARI and NMI values corresponding to the CIC stage, one can infer that the discovered initial communities, in turn, are very high-quality communities that do not require many modifications done by the UP stage. However, as can be seen, the reforms made by the second phase have led to meaningful improvements in communities.

**Table 3:** Comparison results of different stages of TSCDA and also the total time elapsed in the algorithm.

| Network | First stage (CIC) |  |  |  |  |  | Second stage (UC) |  |  |  |  |  |
| --- | --- | --- | --- | --- | --- | --- | --- | --- | --- | --- | --- | --- |
| | $\alpha$ | ARI | NMI | $q_G$ | number of communities | elapsed time (min) | ARI | NMI | $q_G$ | number of iterations | number of communities | elapsed time (min) |
| Karate | 2 | 0.95 | 0.87 | 0.93 | 2 | 0.08 | 1 | 1 | 0.705 | 1 | 2 | 0.02 |
| Dolphin | 5 | 0.56 | 0.51 | 2.4 | 12 | 0.095 | 0.6 | 0.53 | 1.8 | 3 | 8 | 0.065 |
| Football college | 3 | 0.63 | 0.6 | 1.69 | 5 | 0.196 | 0.71 | 0.66 | 0.827 | 3 | 4 | 0.054 |
| Email | 4 | 0.55 | 0.46 | 2.12 | 54 | 1.82 | 0.57 | 0.51 | 1.4 | 8 | 40 | 1.28 |
| Amazon | 15 | 0.51 | 0.448 | 1.17 | 42216 | 76 | 0.54 | 0.49 | 0.72 | 11 | 49146 | 67 |
| Journal | 12 | 0.45 | 0.419 | 1.09 | 258362 | 157 | 0.47 | 0.43 | 0.65 | 9 | 287371 | 137 |
| LFR ($\mu = 0.1$) | 13 | 0.94 | 0.71 | 0.97 | 5 | 0.44 | 0.98 | 0.75 | 0.65 | 6 | 5 | 0.67 |
| LFR ($\mu = 0.3$) | 11 | 0.818 | 0.41 | 1.2 | 9 | 1.1 | 0.85 | 0.54 | 0.89 | 15 | 5 | 1.6 |
| LFR ($\mu = 0.6$) | 6 | 0.427 | 0.417 | 2.21 | 4 | 1.3 | 0.45 | 0.45 | 1.7 | 19 | 4 | 2.08 |
| LFR ($\mu = 0.9$) | 2 | 0.19 | 0.07 | 4.9 | 7 | 1.29 | 0.21 | 0.19 | 3.06 | 25 | 6 | 2.85 |

We conclude this section by noting that our algorithm outperforms the other under-study algorithms in the provided experiments. More specifically, the values of the ARI and NMI obtained by TSCDA is respectively, on average, five and eight percent higher than that of the other algorithms, and according to [56], this small amount of superiority on this scale can lead to significant refinement in the obtained community structures. Furthermore, results in Table 3 clearly show the reliability of initial communities obtained by the CIC stage. It means that we can confidently trust the communities achieved by this stage in extraordinarily massive and dense networks, for which it takes a long time to reform the communities by the UC stage and even apply other community detection algorithms. Therefore, when the network is big enough and the computational time is crucial, while the resulting partitioning quality is not, we can only apply the CIC stage and safely skip the second stage. As a result, we were also succeeded in introducing an almost accurate (but fast) alternative technique to guess relatively high-quality communities in enormous networks.

## 4 Conclusion

This paper presented TSCDA, a community detection algorithm for discovering communities of a given graph. TSCDA used an optimization technique to find communities with respect to optimizing a modified community quality function. The modified function measures communities’ quality based on considering both the compactness and the cohesiveness of communities. The proposed algorithm first establishes appropriate initial communities by estimating proper influential high-degree vertices, which are at a fair distance from each other. Next, it uses a local search technique to achieve final communities with approximately minimum quality values. We propounded six real-world networks and also a category of randomly generated synthetic networks to evaluate TSCDA’s performance against several well-known community detection algorithms. The experimental results show that the ARI and NMI values obtained by our algorithm were approximately, respectively %five and %eight better than the ARI and NMI values obtained by the other algorithms. This superiority in the value of these variables shows that the communities obtained by TSCDA are notably better than the communities obtained by the other algorithms. Based on the results, TSCDA can outperform these algorithms and it can be used as a community detection algorithm to analyze complex networks.

## Footnotes

1 ℂ (*α*) *−*outer degree because the community is not formed yet, so a high degree candidate node must be considered.

2 As mentioned, *α* controls the distance between initial communities, and increasing it will lead the initial influential nodes to be established at a farther distance.

## References

1. Agarwal, G., Kempe, D.: Modularity-maximizing graph communities via mathematical programming. The European Physical Journal B 66(3), 409–418 (2008)

2. Ahmed, F., Abulaish, M.: A generic statistical approach for spam detection in online social networks. Computer Communications 36(10-11), 1120–1129 (2013)

3. Ai, J., Liu, Y., Su, Z., Zhang, H., Zhao, F.: Link prediction in recommender systems based on multi-factor network modeling and community detection. EPL (Europhysics Letters) 126(3), 38003 (2019)

4. Aparicio, S., Villazón-Terrazas, J., Álvarez, G.: A model for scale-free networks: application to twitter. Entropy 17(8), 5848–5867 (2015)

5. Artiles, J., Gonzalo, J., Sekine, S.: The semeval-2007 weps evaluation: Establishing a benchmark for the web people search task. In: Proceedings of the fourth international workshop on semantic evaluations (semeval-2007), pp. 64–69 (2007)

6. Arya, V., Garg, N., Khandekar, R., Meyerson, A., Munagala, K., Pandit, V.: Local search heuristics for k-median and facility location problems. SIAM Journal on computing 33(3), 544–562 (2004)

7. Barabási, A.L., Bonabeau, E.: Scale-free networks. Scientific american 288(5), 60–69 (2003)

8. Blondel, V.D., Guillaume, J.L., Lambiotte, R., Lefebvre, E.: Fast unfolding of communities in large networks. Journal of statistical mechanics: theory and experiment 2008(10), P10008 (2008)

9. Boudebza, S., Cazabet, R., Azouaou, F., Nouali, O.: Olcpm: An online framework for detecting overlapping communities in dynamic social networks. Computer Communications 123, 36–51 (2018)

10. Casciaro, T., Lobo, M.S.: Competent jerks, lovable fools, and the formation of social networks. Harvard business review 83(6), 92–99 (2005)

11. Chakraborty, T., Dalmia, A., Mukherjee, A., Ganguly, N.: Metrics for community analysis: A survey. ACM Computing Surveys (CSUR) 50(4), 54 (2017)

12. Cheikh, S., Sara, B., Sara, Z.: A hybrid heuristic community detection approach. In: 2020 International Conference on INnovations in Intelligent SysTems and Applications (INISTA), pp. 1–7. IEEE (2020)

13. Chen, J., Zäıane, O.R., Goebel, R.: Detecting communities in social networks using max-min modularity. In: Proceedings of the 2009 SIAM international conference on data mining, pp. 978–989. SIAM (2009)

14. Comellas, F., Sampels, M.: Deterministic small-world networks. Physica A: Statistical Mechanics and its Applications 309(1-2), 231–235 (2002)

15. Danon, L., Diaz-Guilera, A., Duch, J., Arenas, A.: Comparing community structure identification. Journal of Statistical Mechanics: Theory and Experiment 2005(09), P09008 (2005)

16. Dao, V.L., Bothorel, C., Lenca, P.: Community structure: A comparative evaluation of community detection methods. Network Science 8(1), 1–41 (2020)

17. Devi, J.C., Poovammal, E.: An analysis of overlapping community detection algorithms in social networks. Procedia Computer Science 89, 349–358 (2016)

18. Ebel, H., Mielsch, L.I., Bornholdt, S.: Scale-free topology of e-mail networks. Physical review E 66(3), 035103 (2002)

19. Ferdowsi, A., Abhari, A.: Generating high-quality synthetic graphs for community detection in social networks. In: Proceedings of the 2020 Spring Simulation Conference, pp. 1–10 (2020)

20. Ferdowsi, A., Khanteymoori, A.: Discovering communities in networks: A linear programming approach using max-min modularity. In: M. Ganzha, L. Maciaszek, M. Paprzycki, D. Ślezak (eds.) Proceedings of the 16th Conference on Computer Science and Intelligence Systems, Annals of Computer Science and Information Systems, vol. 25, p. 329–335. IEEE (2021). DOI 10.15439/2021F65. URL http://dx.doi.org/10.15439/2021F65

21. Flake, G.W., Lawrence, S., Giles, C.L.: Efficient identification of web communities. In: Proceedings of the sixth ACM SIGKDD international conference on Knowledge discovery and data mining, pp. 150–160 (2000)

22. Fortunato, S., Barthelemy, M.: Resolution limit in community detection. Proceedings of the national academy of sciences 104(1), 36–41 (2007)

23. Freitas, L.M., Carneiro, M.G.: Community detection to invariant pattern clustering in images. In: 2019 8th Brazilian Conference on Intelligent Systems (BRACIS), pp. 610– 615. IEEE (2019)

24. Gallier, J.: Spectral theory of unsigned and signed graphs. applications to graph clustering: a survey. arXiv preprint 1601.04692 (2016)

25. Gao, S., Chen, A., Rahmani, A., Jarada, T., Alhajj, R., Demetrick, D., Zeng, J.: Mcf: A tool to find multi-scale community profiles in biological networks. Computer methods and programs in biomedicine 112(3), 665–672 (2013)

26. Ghosh, S., Halappanavar, M., Tumeo, A., Kalyanaraman, A., Lu, H., Chavarria-Miranda, D., Khan, A., Gebremedhin, A.: Distributed louvain algorithm for graph community detection. In: 2018 IEEE international parallel and distributed processing symposium (IPDPS), pp. 885–895. IEEE (2018)

27. Girvan, M., Newman, M.E.: Community structure in social and biological networks. Proceedings of the national academy of sciences 99(12), 7821–7826 (2002)

28. Good, B.H., De Montjoye, Y.A., Clauset, A.: Performance of modularity maximization in practical contexts. Physical Review E 81(4), 046106 (2010)

29. Gupta, A., Tangwongsan, K.: Simpler analyses of local search algorithms for facility location. arXiv preprint 0809.2554 (2008)

30. Hamilton, W.L.: Graph representation learning. Synthesis Lectures on Artifical Intelligence and Machine Learning 14(3), 1–159 (2020)

31. Huang, L., Chao, H.Y., Xie, Q.: Mumod: A micro-unit connection approach for hybrid-order community detection. In: Proceedings of the AAAI Conference on Artificial Intelligence, vol. 34, pp. 107–114 (2020)

32. Hubert, L., Arabie, P.: Comparing partitions. Journal of classification 2(1), 193–218 (1985)

33. Jiang, L., Shi, L., Liu, L., Yao, J., Yousuf, M.A.: User interest community detection on social media using collaborative filtering. Wireless Networks pp. 1–7 (2019)

34. Kim, J., Hastak, M.: Social network analysis: Characteristics of online social networks after a disaster. International Journal of Information Management 38(1), 86–96 (2018)

35. Kumar, S., Hanot, R.: Community detection algorithms in complex networks: A survey. In: International Symposium on Signal Processing and Intelligent Recognition Systems, pp. 202–215. Springer (2020)

36. Lancichinetti, A., Fortunato, S., Radicchi, F.: Benchmark graphs for testing community detection algorithms. Physical review E 78(4), 046110 (2008)

37. Leskovec, J., Kleinberg, J., Faloutsos, C.: Graph evolution: Densification and shrinking diameters. ACM Transactions on Knowledge Discovery from Data (TKDD) 1(1), 2 (2007)

38. Linares, O.A., Botelho, G.M., Rodrigues, F.A., Neto, J.B.: Segmentation of large images based on super-pixels and community detection in graphs. IET Image Processing 11(12), 1219–1228 (2017)

39. Lusseau, D., Schneider, K., Boisseau, O.J., Haase, P., Slooten, E., Dawson, S.M.: The bottlenose dolphin community of doubtful sound features a large proportion of long-lasting associations. Behavioral Ecology and Sociobiology 54(4), 396–405 (2003)

40. Malliaros, F.D., Vazirgiannis, M.: Clustering and community detection in directed networks: A survey. Physics Reports 533(4), 95–142 (2013)

41. Mkhitaryan, K., Mothe, J., Haroutunian, M.: Detecting communities from networks: Comparison of algorithms on real and synthetic networks

42. Newman, M.E.: Fast algorithm for detecting community structure in networks. Physical review E 69(6), 066133 (2004)

43. Newman, M.E.: Modularity and community structure in networks. Proceedings of the national academy of sciences 103(23), 8577–8582 (2006)

44. Newman, M.E., Barabási, A.L.E., Watts, D.J.: The structure and dynamics of networks. Princeton university press (2006)

45. Orman, G.K., Labatut, V.: A comparison of community detection algorithms on artificial networks. In: International conference on discovery science, pp. 242–256. Springer (2009)

46. Perliger, A., Pedahzur, A.: Social network analysis in the study of terrorism and political violence. PS: Political Science & Politics 44(1), 45–50 (2011)

47. Radicchi, F., Castellano, C., Cecconi, F., Loreto, V., Parisi, D.: Defining and identifying communities in networks. Proceedings of the National Academy of Sciences 101(9), 2658–2663 (2004)

48. Raghavan, U.N., Albert, R., Kumara, S.: Near linear time algorithm to detect community structures in large-scale networks. Physical review E 76(3), 036106 (2007)

49. Rosvall, M., Bergstrom, C.T.: Maps of random walks on complex networks reveal community structure. Proceedings of the National Academy of Sciences 105(4), 1118–1123 (2008)

50. Schaeffer, S.E.: Graph clustering. Computer science review 1(1), 27–64 (2007)

51. Shang, R., Liu, H., Jiao, L., Esfahani, A.M.G.: Community mining using three closely joint techniques based on community mutual membership and refinement strategy. Applied Soft Computing 61, 1060–1073 (2017)

52. Shchur, O., Günnemann, S.: Overlapping community detection with graph neural networks. arXiv preprint 1909.12201 (2019)

53. Shen, H.W., Cheng, X.Q., Guo, J.F.: Quantifying and identifying the overlapping community structure in networks. Journal of Statistical Mechanics: Theory and Experiment 2009(07), P07042 (2009)

54. Shi, J., Malik, J.: Normalized cuts and image segmentation. Departmental Papers (CIS) p. 107 (2000)

55. Shi, X., Lu, H., He, Y., He, S.: Community detection in social network with pairwisely constrained symmetric non-negative matrix factorization. In: 2015 IEEE/ACM International Conference on Advances in Social Networks Analysis and Mining (ASONAM), pp. 541–546. IEEE (2015)

56. Sobolevsky, S., Campari, R., Belyi, A., Ratti, C.: General optimization technique for high-quality community detection in complex networks. Physical Review E 90(1), 012811 (2014)

57. Varsha, K., Patil, K.K.: An overview of community detection algorithms in social networks. In: 2020 International Conference on Inventive Computation Technologies (ICICT), pp. 121–126. IEEE (2020)

58. Von Luxburg, U.: A tutorial on spectral clustering. Statistics and computing 17(4), 395–416 (2007)

59. Wang, M., Wang, C., Yu, J.X., Zhang, J.: Community detection in social networks: an in-depth benchmarking study with a procedure-oriented framework. Proceedings of the VLDB Endowment 8(10), 998–1009 (2015)

60. Wei, Y.C., Cheng, C.K.: Towards efficient hierarchical designs by ratio cut partitioning. In: 1989 IEEE International Conference on Computer-Aided Design. Digest of Technical Papers, pp. 298–301. IEEE (1989)

61. Wenping, Z., Chenhao, C., Yuhua, Q., Jie, W.: A two-stage community detection algorithm based on label propagation. Journal of Computer Research and Development 55(9), 1959 (2018)

62. Whang, J.J., Gleich, D.F., Dhillon, I.S.: Overlapping community detection using neighborhood-inflated seed expansion. IEEE Transactions on Knowledge and Data Engineering 28(5), 1272–1284 (2016)

63. Xie, J.R., Wang, B.H.: Modularity-like objective function in annotated networks. Frontiers of Physics 12(6), 128903 (2017)

64. Xu, G., Guo, J., Yang, P.: Tns-lpa: An improved label propagation algorithm for community detection based on two-level neighbourhood similarity. IEEE Access 9, 23526–23536 (2020)

65. Yang, J., Leskovec, J.: Defining and evaluating network communities based on groundtruth. Knowledge and Information Systems 42(1), 181–213 (2015)

66. Yip, K.Y., Cheung, D.W., Ng, M.K.: Harp: A practical projected clustering algorithm. IEEE Transactions on knowledge and data engineering 16(11), 1387–1397 (2004)

67. Zachary, W.W.: An information flow model for conflict and fission in small groups. Journal of anthropological research 33(4), 452–473 (1977)

68. Zhang, W., Shang, R., Jiao, L.: Complex network graph embedding method based on shortest path and moea/d for community detection. Applied Soft Computing 97, 106764 (2020)

